# Unique Adaptations of a Photosynthetic Microbe *Rhodopseudomonas palustris* to the Toxicological Effects of Perfluorooctanoic Acid

**DOI:** 10.1101/2025.02.19.638955

**Authors:** Mark Kathol, Anika Azme, Sumaiya Saifur, Nirupam Aich, Rajib Saha

**Author notes:** These authors have contributed equally to this work and share the first authorship (Mark Kathol, Anika Azme).

## Abstract

In this study, we investigate the PFOA removal capabilities of *Rhodopseudomonas palustris* (*R. palustris*), a fluoroacetate dehalogenase containing microbe as a potential candidate for achieving bioremediation. In the 50-day PFOA uptake experiment, R. palustris removed 44 ± 6.34 % PFOA after 20 days of incubation, which was then reduced to a final removal of 6.23 ± 12.75 %. Results indicate PFOA was temporarily incorporated into the cell membrane before being released partially into the media after cell lysis. This incorporation might be attributed to the combined effect of hydrophobic interaction between PFOA and the cell membrane and the reduced electrostatic repulsion from the high ion presence in the growth medium. The growth of *R. palustris* during the PFOA uptake experiment was 9-fold slower than their growth without PFOA. This study also completely defines the toxicity range of PFOA for *R. palustris* through a toxicity assay. Increasing PFOA concentration reduced the microbe growth, with complete inhibition around 200 ppm. For various concentrations of PFOA, R. palustris exhibits interesting diauxic growth behavior. An accelerated growth phase was followed by a temporary death phase in the first 24 hours in the presence of 12.5-100 ppm PFOA, implying a unique adaptation mechanism to PFOA.

## INTRODUCTION

Per- and polyfluoroalkyl substances (PFAS) are a group of >10,000 fluorinated synthetic compounds that are highly toxic and environmentally persistent. These compounds are difficult to break down due to the high energy C-F bonds (Son et al., 2020). This leaves a large PFAS accumulation in our soil and water resources and with concerns for potential human and ecological toxicity (Sinclair et al., 2020; Wang et al., 2023). PFAS has shown toxicological properties against environmental soil microbes including negative effects on soil respiration and soil bacteria enzyme activity (Xu et al., 2023). Also, when a soil microbial community was exposed to perfluorooctanoic acid (PFOA; a priority PFAS), the growth of soil microbes were reduced, however, the negative effects are somewhat temporary and the community recovered within a year (Xu et al., 2022).

Conversely, microbial degradation of PFAS is considered one of the most desired PFAS remediation strategies due to the advantages of lower capital and operational costs than abiotic methods (Shahsavari et al., 2021; Zhang et al., 2022). Many recent studies have attempted to find organisms from high PFAS concentration effluent streams that utilize PFAS as a potential carbon source for growth (Marciesky et al., 2023). However, due to strong C-F bonds, PFAS are not nearly as attractive for bacteria to utilize as a carbon source compared to others, such as sugars or lignocellulosic biomass, and there is no known enzymatic method to utilize it as an energy source (Harper et al., 2003). However, several “proof-of-concept” studies have demonstrated the potential of certain microbial species to degrade PFAS. These include *Acidimicrobium* sp. strain A6 (Ruiz-Urigüen et al., 2022), dehalogenase possessing strains such as *Dechloromonas* sp. CZR5 (Long et al., 2024), and various *Pseudomonas* species, such as *Pseudomonas parafulva* (Yi et al., 2019; Yi et al., 2016).

*Rhodopseudomonas palustris* (Hereafter, *R. palustris*) is a highly adaptive gram-negative, non-sulfur bacterium that can be found in diverse environmental media, such as soil, aquatic sediment, eutrophic ponds (Rayyan et al., 2018), animal waste lagoons (Kim et al., 2004), and wastewater (Harwood, 2022). *R. palustris* is known for its high environmental hardiness, resisting the toxicological effects of high metal concentrations, and even removing them under these conditions (Llorens et al., 2012; Zhao et al., 2015). Furthermore, *R. palustris* is noteworthy for its ability to break down recalcitrant carbon sources such as lignin breakdown products (Alsiyabi et al., 2021; Brown et al., 2022; Harwood and Gibson, 1988). This strain also possesses a fluoroacetate dehalogenase (*rpa1163*) enzyme. When purified from bacteria (not *R. palustris*), this enzyme has been observed to cleave the very strong C−F bond from fluoroacetate (Chan et al., 2010) and 2,3,3,3-tetrafluoropropionic acid (Li et al., 2019). To assess the potential of *R. palustris* for PFAS biodegradation, it is crucial to study its interactions with PFAS and directly evaluate the toxic effects of these compounds on its morphology and metabolic processes. Therefore, this study aims to pave the road for future PFAS bioremediation studies involving *R. palustris* and gram-negative bacteria in general by characterizing *R. palustris’* growth and adaptive capabilities when exposed to PFOA as well as PFOA uptake and toxicity to this organism.

## 2. METHODOLOGY

To achieve our above-mentioned objectives, first, we performed a 50-day growth study for *R. palustris* in a photosynthetic media (PM) with exposure to 50 ppm PFOA in anaerobic conditions. The toxicity effects of PFOA were evaluated by monitoring time dependent bacterial growth through optical density measurements at 660 nm (OD_660_) and by evaluating physical cell damage via transmission electron microscopy (TEM) imaging. The PFOA uptake by *R. palustris* was evaluated by measuring changes in PFOA concentration in PM at different time intervals using liquid chromatography coupled with tandem mass spectrometry (LC-MS/MS). The underlying mechanisms were further explored via surface charge measurements of *R. palustris* cells using a Zetasizer and anion concentration analysis in PM using ion chromatography (IC). Moreover, a 5-day dose-dependent toxicity assay was performed using PM spiked with PFOA concentrations ranging from ∼0.78 to 200 ppm, and *R. palustris* growth was measured using OD_660_. Detailed descriptions of methodology are provided in the supplementary information (SI, Section 1).

## 3. RESULTS AND DISCUSSION

### 3.1. PFOA Uptake by *R. palustris*

Figure 1 presents the trend of PFOA uptake from the PFOA-spiked PM containing lysed (control) and live (sample) cell cultures. The PFOA concentration on day 0 was less than the desired 50 ppm initial concentration for both the live samples (36.14 ± 5.00 ppm) and lysed control groups (23.99 ± 2.57 ppm). A prior check of 50 ppm PFOA spiked PM without any culture reliably returned concentrations of near 50 ppm, and PFOA from the reactor did not show any degradation products. Therefore, the discrepancy is attributed to the combination of the complex mixture (culture presence in PM) and the potential adsorption on glass container used for the anaerobic reactor. The PFOA removal for each group (live or lysed) was calculated using the PFOA concentration in day 0 in respective samples (Eq. S1). In the live sample group, the PFOA removal percentage first gradually increased to 44 ± 6.34 % until day 20, stabilized from day 20-30, and then showed significant decrease to 6.23 ± 12.75 % at day 50. In the lysed group, PFOA removal reached 12.19 ± 16.30 % on day 20, but final concentrations at 50 days showed no uptake by the lysed cells. A similar observation has been reported for two *Pseudomonas* sp. strains, where Perfluorohexane Sulfonate (PFHxS) release happened after 15 days of incubation (Presentato et al., 2020).

**Figure 1.**
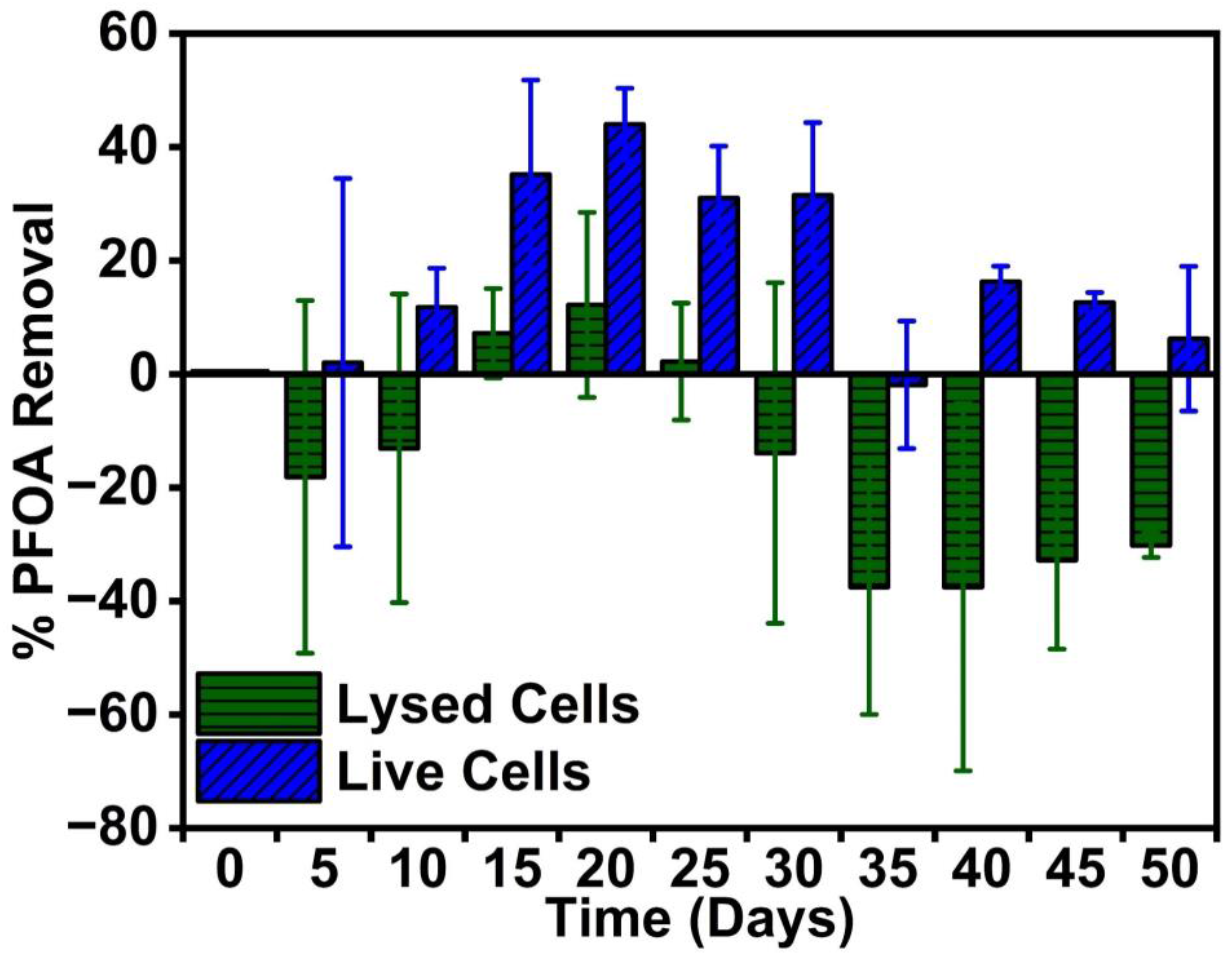
PFOA removal trends in lysed and live *R. palustris* cells grown in a 50 ppm PFOA spiked PM over 50 days. Error bars represent one standard deviation for n = 3 samples.

The significant difference in the PFOA removal percentage between the live and lysed bacteria indicates that more PFOA is uptaken by live *R. palustris* cells lysed bacteria. The gradual increase in PFOA uptake (until day 20) and release in the later days might be a result of a complex interaction process that include physical hydrophobic and electrostatic interactions along with biological ion transport mechanisms. The uptaken PFOA might be incorporated into the phospholipid bilayer of the cell membrane, likely due to hydrophobic interaction (Fitzgerald et al., 2018). The PFOA decrease in PM might be due to the physical PFOA adsorption on the outside cell surface and the incorporation inside the cell. When suspended in deionized (DI) water, the live and lysed *R. palustris* cells from day 20 have shown high negative charges of –36.9 ± 0.977 mV and –43.9 ± 0.912 mV, respectively (Figure S2.1). Both PFOA and *R. palustris* have high negative surface charges, so electrostatic repulsion would hinder the adsorption of PFOA on the cells. However, the high ion concentrations (calculated to be 189.42 mM) in the PFOA-spiked PM reduced the surface charges to –9.31 ± 0.83 mV and –14.2 ± 0.625 mV for the live and lysed cell cultures, respectively (Figure S2.1). This would allow PFOA to be near the cells and partition to cell membranes driven by hydrophobicity. However, throughout the 50-day experiment, live bacteria samples uptake more PFOA than lysed cells. Because a higher negative surface charge on the lysed bacteria would result in stronger electrostatic repulsion between PFOA and the lysed cell than the live bacteria. Furthermore, while active ion transport in live bacteria might facilitate PFOA uptake, the absence of functional ion transport mechanisms in lysed cells prevented PFOA uptake. Moreover, due to the inactive ion transport and damaged cell membranes, the charged molecules might leak from within the lysed cell and accumulate on the cell surface (Ayala-Torres et al., 2014). This would explain the lysed cells having more negative surface charge than live bacteria, like other studies (Ayala-Torres et al., 2014; Martinez et al., 2008). Furthermore, bacterial growth in the live group might increase biomass content over time compared to the lysed group, contributing to higher PFOA removal by live bacteria than by lysed ones. Cell lysis might be responsible for the eventual partial release of the incorporated PFOA (Marchetto et al., 2021).

### 3.2. Free Anion Mass Balance in PM

Since day 20 exhibited the highest PFOA removal (Figure 1), the live group PM was further analyzed to determine possible nutrient consumption and identify other anion generation, such as fluoride (F^-^). IC results showed 90.01 ± 7.64 ppm chloride, 3051.30 ± 242.62 ppm phosphate, and 2279.56 ± 214.84 ppm sulfate in the PM at day 20 (Figure S2.2). While the chloride and sulfate concentrations remained constant in the PM, the phosphate concentration decreased 67.87% from the initial concentration of 9497 ppm. As phosphate acts as a crucial nutrient for the cell development and function of *R. palustris*, this decrease in phosphate amount in 20 days indicated the growth of *R. palustris* in PFOA spiked PM (Berne et al., 2005; Kim et al., 2004). However, no fluoride ion was detected in the PM, indicating that the PFOA uptaken by the microbe had only been stored in the cells and not metabolized.

### 3.3. *R. palustris* Growth Under PFOA Exposure

To observe the effects of PFOA on the growth of *R. palustris*, OD_660_ was measured every sampling day (Figure S2.3a). Ideally, this would have produced a curve where the growth of *R. palustris* cultures exposed to PFOA would either be identical to, or slightly higher than, the control acetate curves. However, wild type *R. palustris*, when incubated anaerobically in the 250 mL system, grows extremely slower than our acetate control growth curves. An anaerobic wild type *R. palustris* culture would typically take 3-5 days to reach a maximum OD_660_ of roughly 0.6 (Figure S2.3b) (Brown et al., 2022; Brown et al., 2020; Immethun et al., 2022), whereas our 50 ppm PFOA-exposed live cultures grew at an incredibly slow rate, taking 45 days until it reached the same maximum OD_660_. This, because in our PFOA-exposed cultures, 20 mM of sodium acetate was also supplemented, strongly suggests that our cultures are continuously experiencing cell lysis from the toxicological effects of PFOA. A second noticeable trend we observed was an initial, large, albeit temporary, increase in OD_660_ and the presence of two growth phases, or diauxic growth (before and after day 10). This type of growth usually occurs in cultures with two carbon sources, where an organism preferentially consumes one over the other (Mahadevan et al., 2002; Sun et al., 2020; Tyrovouzis et al., 2014). However, due to the lack of fluoride present in the media after growth and the inconsistent PFOA mass balance, this may instead be the result of an adaptive phase, where *R. palustris* responds to PFOA exposure. The apparent accelerated death phase could then be explained by cell lysis before *R. palustris* fully adapts. pH change was also trivial for the duration of the experiment, owing to the high amount of phosphate present in the media (Figure S2.4).

### 3.4. PFOA Effect on *R. palustris* Morphology

To further evaluate the toxic effects of PFOA, the TEM images (Figure S2.5) provided additional information on the live cell structure and growth in the presence and absence of PFOA. Healthy wild type *R. palustris* cell cultures without PFOA exposure were observed to contain nucleoid and smooth cell membranes (Figure S2.5 a & b). However, the high magnification images showed PFOA-exposed live cells from days 0, 20, and 35 with visible damages from possible oxidative stress, such as the absence of nucleoids, cytoplasm leakage, cell wall disruption, and uneven cell membranes (Figure S2.5 d, f, & h). Similar damages in different bacterial cells from PFAS exposure have been reported in other works (Lindell et al., 2024; Presentato et al., 2020). Even with these toxic effects, the low-magnification images (Figure S2.5 b, e, & g) showed an increasing number of live cells from day 0 to 35 under PFOA exposure. This corroborated with the growth curve (Figure S2.3), indicating the ability of *R. palustris* to grow in the presence of a high concentration of PFOA (i.e., 50 ppm).

### 3.5. Dose-dependent Toxicity Assay

Given that our initial 50 ppm experiment produced very irregular growth and morphological damage, A 5-day dose-dependent toxicity assay was performed where wild type *R. palustris* was incubated with a gradient of PFOA concentrations. The purpose of this experiment was to reproduce the irregular growth profile observed in the 50 ppm concentration cultures to determine if this effect also exists, or is amplified, for other concentrations of PFOA. A 2x dilution of PFOA to add to wild type *R. palustris* cultures ranging from final concentrations of 200 ppm to ∼0.78 ppm, with controls containing no PFOA (Figure 2). From our results, two very interesting trends emerge; firstly, the initial large temporary increase in OD_660_, which occurs in the first 24 hours, is present for PFOA concentrations above 12.5 ppm. This effect begins at 12.5 ppm and then increases in intensity until 100 ppm, where higher concentrations almost completely inhibit sustained *R. palustris* growth. Secondly, this may seem to emulate diauxic growth behavior, however, considering the lack of PFOA degradation as seen from our mass balance and our IC results, we hypothesize that this may be a period where *R. palustris* responds to the toxicological effects of PFOA. However, to the best of our knowledge, we have not seen any literature that exhibits a similar growth pattern to the one we have obtained. This could imply a unique resistance or adaptation mechanism that *R. palustris* possesses, allowing it to grow even in very high toxin concentrations. A recent study demonstrated the ability of PFOA to incorporate itself into lipid bilayers up to a saturation limit, expanding it and increasing membrane fluidity (Sobolewski et al., 2024). We hypothesize that the toxicity gradient we observed results from this phenomenon, where PFOA is saturating the cell membrane until it is ruptured. However, the underlying mechanism of R. palustris to effectively respond to and recover from this toxicological effect remains to be understood.

**Figure 2.**
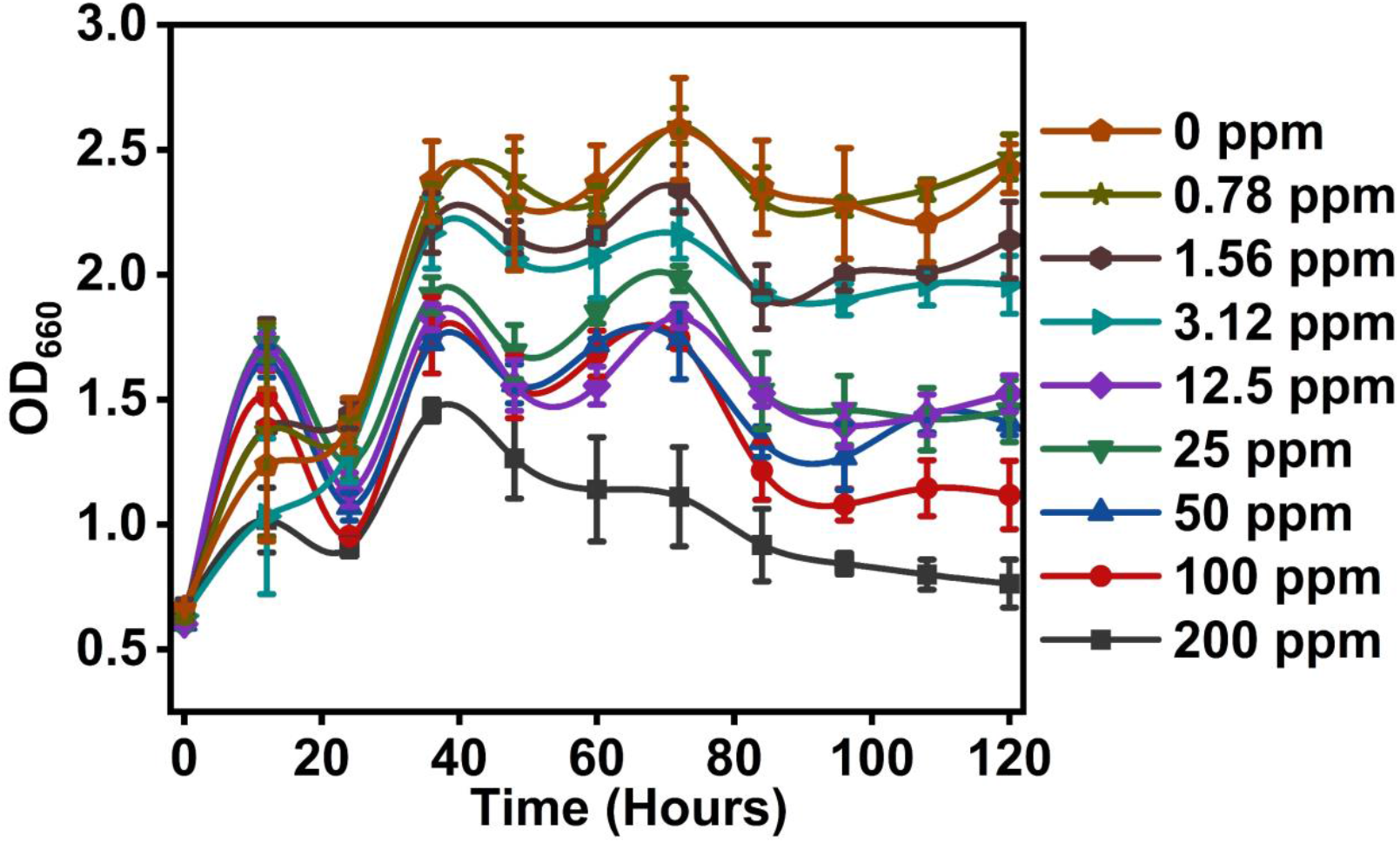
Wild type *R. palustris* grown on a 2x serial dilution of PFOA (200 - 0.78 ppm) over 5 days. The growth curves showed an accelerated death phase in the initial hours (12-24 hours) for the PFOA concentrations of 12.5-200 ppm. With the increase in PFOA concentration, the final OD_660_ decreased, and growth stopped at 200 ppm PFOA. Error bars represent one standard deviation for n = 3 samples.

## 4. CONCLUSIONS

In summary, this study presents the potential of *R. palustris* for PFOA removal and elucidates the microbes’ unique adaptation pattern to the toxic effects of PFOA. At high PFOA concentration of 50 ppm, *R. palustris* temporarily removed PFOA from the surrounding media either by incorporation into or adsorption on the cell membrane. While the PFOA removal reached a maximum uptake of ∼44% after 20 days, eventual PFOA release happens from the cell membrane. The full toxicity range of PFOA on *R. palustris* is determined, where cultures exposed to ∼0.78 ppm of PFOA are identical to cultures grown without PFOA. The growth inhibition increases with increasing PFOA concentration and is completely inhibited at 200 ppm. *R. palustris* also exhibits interesting growth adaptation behavior, where an initial growth phase is followed by an accelerated death phase due to the high concentration of PFOA. Another growth phase follows when *R. palustris* cultures are supplemented with 12.5 ppm - 100 ppm PFOA. The dehalogenase enzymes *R. palustris* possesses might function at lower concentrations of other halogenated compounds (Li et al., 2021). However, this study demonstrates that a PFOA concentration ceiling limit exists for effective bioremediation; beyond that, *R. palustris* suffers severe toxicity effects. Moreover, the PFOA adsorption effects of either the cell membrane or the vessel surfaces, or a combination of both, make direct measurement of the PFOA concentration in the media difficult. Accurate mass balancing is crucial to determining PFOA removal; therefore, we recommend that future bioremediation studies use additional controls to verify the consumption of PFOA.

## ACKNOWLEDGMENT

This work was supported by the University of Nebraska-Lincoln Layman Seed Grant and Nebraska Collaboration Initiative Grant awarded to R.S. and N.A. This work was also supported by the National Science Foundation CAREER Grant (1943310) awarded to R.S. This work was partially supported by National Science Foundation Grant (2324853) awarded to N.A. The research was partly performed at the UNL Nebraska Center for Biotechnology. We thank Dr. You Zhou and Dr. Bara Altartouri from the NCB Microscopy Core Research Facility for their assistance with the TEM sample preparation and imaging.

