## Supplementary File for "Unique Adaptations of a Photosynthetic Microbe *Rhodopseudomonas palustris* to the Toxicological Effects of Perfluorooctanoic Acid"

### S1. Materials and Methods

#### S1.1. Strain Growth Conditions

Wild-type *R. palustris* (Molisch) van Niel BAA-98 CGA009 was obtained from the American Type Culture Collection (ATCC). When not in use, strains were stored at -80°C at a final concentration of 20% glycerol in sterile freezer tubes. When retrieved from storage, *R. palustris* was first streaked onto solid 112 Van Niel's (VN) media. Seed cultures of *R. palustris* were grown aerobically before performing growth experiments in Photosynthetic media (PM), prepared as described previously (Kathol et al., 2023). These seed cultures were grown in 250 mL Erlenmeyer flasks with 50 mL of PM, prepared as described previously (Kathol et al., 2023). Seed cultures were then supplemented with 20 mM sodium acetate, 15.2 mM $(NH_4)_2SO_4$, and 10 mM $NaHCO_3$. All cultures of *R. palustris* were grown at 30°C and 275 rpm.

#### S1.2. Culture Preparation and PFOA Uptake Experiment

250 mL of triplicate anaerobic *R. palustris* cultures were prepared by diluting seed cultures to $OD_{660}$ = 0.2 in 500 mL glass Erlenmeyer flasks supplemented with 20 mM sodium acetate ($CH_3COONa$), 15.2 mM $(NH_4)_2SO_4$, and 10 mM $NaHCO_3$ in PFOA-containing PM, and stoppered with sample collection and purge tubes. The perfluorooctanoic acid, PFOA ($CF_3(CF_2)_6COOH$), CAS 335-67-1, molecular weight 414.07, 100%) was purchased from Toronto Research Chemicals, Canada. 50 ppm PFOA was prepared by the direct addition of 50 mg PFOA to 1 L PM in polypropylene bottles. To ensure the correct addition of PFOA to a final concentration of 50 ppm, cell cultures were removed from regular PM by centrifugation in a Beckman Coulter Allegra X-30R Centrifuge (Beckman Coulter Life Sciences, Indiana, USA) at 3500 × g for 15 minutes in sterile 50 mL Falcon Tubes. Cells were resuspended in PFOA-containing PM. The 50-day growth experiment consisted of a triplicate control group (lysed cells) through autoclaving at 121°C for

45 minutes, and a sample group containing live cells. Both groups were prepared by dilution from seed cultures to $OD_{660}$ = 0.2. Both lysed and live cultures were then incubated with an Algaetron PSI incubator under 100 μE of light at 30°C and 275 rpm.

10 mL aliquots were taken from both the lysed and live cultures every 5 days, including day 0, and *R. palustris* cultures were also re-supplemented with 2 mM sodium acetate to ensure continuous growth over the 50 days. $OD_{660}$ of each aliquot was measured using a Genesis 10S UV-Vis Spectrophotometer, and the pH was then measured using an accumet$^{TM}$ pH probe. All aliquots were then immediately stored at -80°C. These aliquotes were used for PFOA quantification using an ultrahigh pressure liquid chromatography (UPLC) system interfaced with a triple quadrupole mass spectrometer system (Waters Corporation, Milford, MA). The day 20 aliquots were analyzed using IC to measure the anion concentration change in PM. Detailed sample preparation and analysis using LC-MS/MS and IC are discussed below.

The live cell suspensions from days 0, 20, and 35 were returned to room temperature and centrifuged at 4000 rpm for 20 minutes to prepare solid pellets of the *R. palustris* cells for TEM imaging to determine the physical damage. The day 20 live and lysed cells were used for measuring the surface charge in deionized water as well as in PM media with and without PFOA using a Malvern ZetaSizer Nano ZS90 instrument (details below).

**S1.3. Dose-dependent Toxicity Assay**

Growth curves for the toxicity assay were obtained using sterilized polypropylene 48-well plates. Seed cultures were diluted similarly to a final concentration of $OD_{660}$ = 0.2 and supplemented with 20 mM $CH_3COONa$, 15.2 mM $(NH_4)_2SO_4$, and 10 mM $NaHCO_3$ in PM. *R. palustris* well-cultures were prepared in triplicate with PFOA concentrations in a 2x serial dilution range from 200 ppm - ~0.78 ppm ($200 - 200\times2^{-8}$ ppm). Air was purged from the plate using nitrogen, and resazurin dye was used to ensure anaerobic conditions (Wagner et al., 2019). The plate was then caulked shut and incubated at 30°C and 275 rpm for 5 days. $OD_{660}$ was measured every 12 hours, including hour 0, using a plate reader with an area scan protocol.

**S1.4. Physical Cell Damage Characterization**

After removing the supernatant, 2 mL fixation solution glutaraldehyde/ paraformaldehyde (2% each in 0.1 M cacodylate buffer) was added slowly to the centrifugation tube, ensuring no pellet dislodgement. The mixed solution was left undisturbed for 45 minutes to allow it to set. The sample was also post-fixed with 1% osmium tetroxide for an hour at room temperature. The sections for the TEM grid were cut with an ultramicrotome using a diamond blade (DiATOME knife), and the thickness of the ultrathin sections was around 75 nm. An advanced Transmission Electron Microscope, Hitachi HT-7800 TEM (Santa Clara, CA, USA), with an accelerating voltage of 80 kV, placed at the Nebraska Center for Biotechnology at the University of Nebraska-Lincoln (UNL), was used to capture images of control wild-type and PFOA-spiked *R. palustris* live cells. Images were captured at 5 μm and 500 nm magnifications to determine the physical cell damage from PFOA exposure.

**S.1.5. Surface Charge Measurement**

The Malvern ZetaSizer Nano ZS90, equipped with a 4 mW He−Ne 633 nm laser (Malvern Instrument, Worcestershire, UK), was used to measure the surface charge of the *R. palustris* cells from day 20. The optical density of the live and lysed cells used for this measurement was 0.23 (measured at 660 nm). The cells were dispersed in water, PM, and PFOA-spiked PM media. The zeta potential was measured by placing 850 μL of live or lysed cell dispersion in the disposable Malvern Panalytical folded capillary zeta cell. At least 10 electrophoretic mobility measurements were recorded for each dispersion.

##### S1.6. Quantitative Analysis

###### S1.6.1. Liquid Chromatography-Mass Spectrometry (LC-MS/MS) Analysis

The stored aliquots were allowed to reach room temperature before processing. The vials were vortexed before two-step dilution was performed to reduce the concentration of PFOA from 50 ppm to 125 ppb. 12.5 µL of these diluted solutions were added to a 15 mL polypropylene centrifuge tube, followed by 100 µL of internal standard, 100 µL of surrogate spike, and methanol, for a total volume of 312.5 µL. The final composition contained 2 ng of internal standard and 4 ng of surrogate spike each, with a methanol-water ratio of 96:4. This solution was then vortexed for 15 seconds before being filtered using a 5 mL syringe with a 25 mm 0.45 µm nylon fiber filter. This step was performed to eliminate any residual biological material from the reactor system. The filtered sample was transferred to a 300 µL polypropylene screw-neck vial for the analysis. For each analysis, quality assurance and quality control samples were prepared similarly. All samples were stored in a -20°C freezer until ready to run.

The peak integration and analysis of concentrations from the standard curve were performed using the Waters MassLynx Software v4.2. For each quantification, a total of seven standards ranging from 0- 20 ng/ mL were prepared for the calibration curve of the instrument. The mobile phase composition was selected from “A” (2 mM ammonium acetate in water) and “B” (2 mM ammonium acetate in methanol). Initial mobile conditions were 95:5 A/B followed by a linear gradient to obtain a composition of 75:25 A/B at 0.5 min, then increased to 50:50 A/B at 3 min, 15:85 A/B at 6 min, and 5:95 A/B at 6.5 min. This was then held for 9 minutes before returning to the initial conditions. The total run time was 12 minutes per sample. A VanGuard pre-column (5 mm × 2.1 mm) was used as a guard column, an isolator column (2.1 mm × 50 mm), and a 50 µL sample extension loop. The UPLC system was equipped with a BEH C18 reverse phase HPLC column (50 mm × 2.1 mm × 1.7 μm). The mass spectrometer used a Unispray™ system operating on negative ion detection mode. The method detection limit of the water sample was calculated to be 0.1 μg/ L. The instrument detection limit for the direct injection methods ranged from 0.034 to 0.245 μg/ L on-column for the lowest calibration standard.

**S.1.6.2. Ion Chromatography Analysis**

2 mL of undiluted aliquots collected on Day 20 were filtered using 5 mL plastic syringes with clean 25 mm 0.45 μL polyethersulfone membrane filters and placed in 1.5 mL autosampler vials. The same aliquots were analyzed at 40x and 400x dilutions using a freshly prepared eluent solution (3.5 mM $Na_2CO_3$ and 1 mM $NaHCO_3$) to detect anion peaks. Lab reagent blanks were included for each analysis to measure possible contamination from reagents, equipment, or the environment. Lab-fortified blanks with a spiked concentration of 3 ppm of each anion were also added to measure the accuracy and recovery efficiency of the analytical method.

The possible fluoride formation from PFAS degradation was analyzed using the Thermo Dionex ICS 5000+ Ion Chromatography System equipped with a Dionex IonPac AS14 4×250 mm column. The EPA method 300.0 was followed with a slight adjustment of 3.5 mM $Na_2CO_3$ and 1 mM $NaHCO_3$ as the eluent solution. Seven standards ranging from 0 to 10 mg/L were prepared for the calibration curve. Using this method, the accepted fluoride, chloride, orthophosphate, and sulfate concentration ranges were 0.0096- 410 mg/ L, 0.0238- 410 mg/ L, 0.0517- 410 mg/L, and 0.0606- 410 mg/L, respectively. The retention times of fluoride, chloride, orthophosphate, and sulfate were 2.4, 3.5, 8.1, and 10 min, respectively. The peak integration and analysis of concentrations from the standard curve were performed using the Chromeleon 7 software.

##### S1.7. PFOA Removal Calculation

The concentration of PFOA was determined using the following equation:

$$Removal\ (\%) = \frac{C_0 - C_x}{C_0} \times 100 \text{ ......... Eq. S1}$$

Where $C_0$ = PFOA concentration in PM on Day 0

$C_x$ = PFOA concentration in PM on Day x (x = 5, 10, 15, …, 50)

#### S2. Results

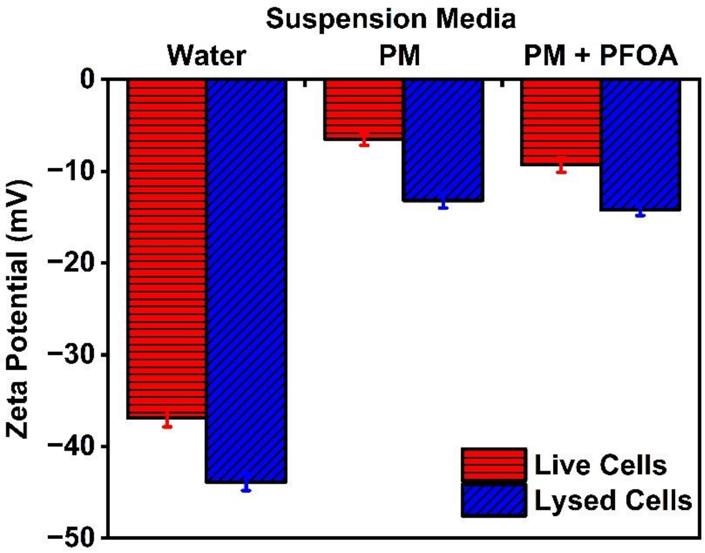


**Figure S2.1:** Surface charge of live and lysed *R. palustris* suspended in water, PM, and PFOA-spiked PM. Live cells have decreased surface charge compared to lysed cells in all suspension media. The presence of ions in PM decreases the surface charges of live and lysed cells suspended in PM. Error bars represent one standard deviation for n = 3 samples.

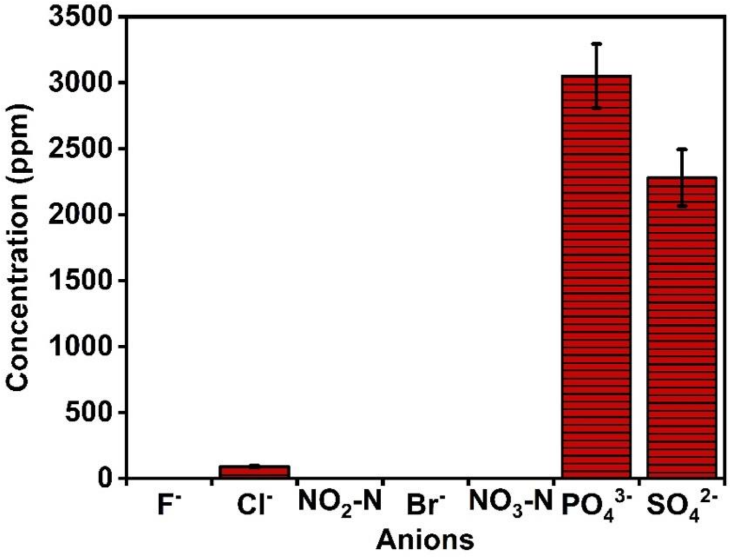


**Figure S2.2:** Anions concentration present in the Day 20 sample group. High chloride (90.01 ± 7.64 ppm), orthophosphate (3051.30 ± 242.62 ppm), and sulfate (2279.56 ± 214.84 ppm) anion concentrations were present in the sample originating from the PM media. No fluoride, nitrite, nitrate, or bromide was detected in the PM. Error bars represent one standard deviation for n = 3 samples.

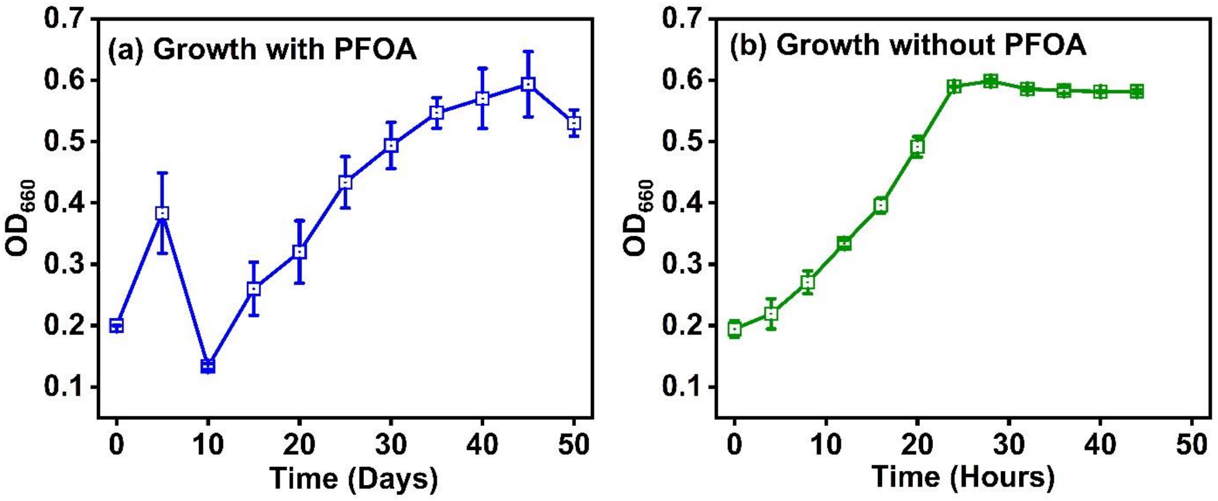


**Figure S2.3:** Growth of *R. palustris* in (a) presence and (b) absence of PFOA-spiked PM. *R. palustri*s reached $OD_{660}$ = 0.6 at around 45 days in the presence of PFOA, compared to 24 hours in its absence. The growth curve in PFOA presence also showed a unique death phase in the initial incubation period (Days 5-10). Error bars represent one standard deviation for n = 3 samples.

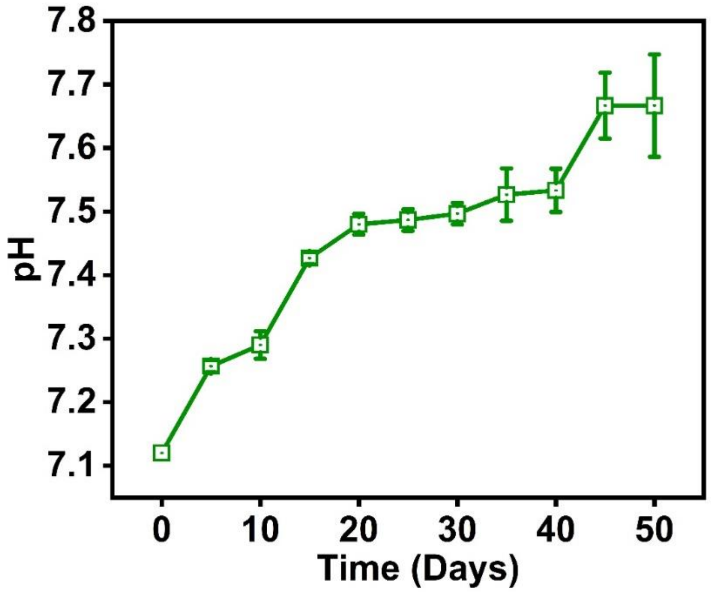


**Figure S2.4:** pH change over the 50 day period for the PFOA uptake experiment. Error bars represent one standard deviation for n = 3 samples.

161

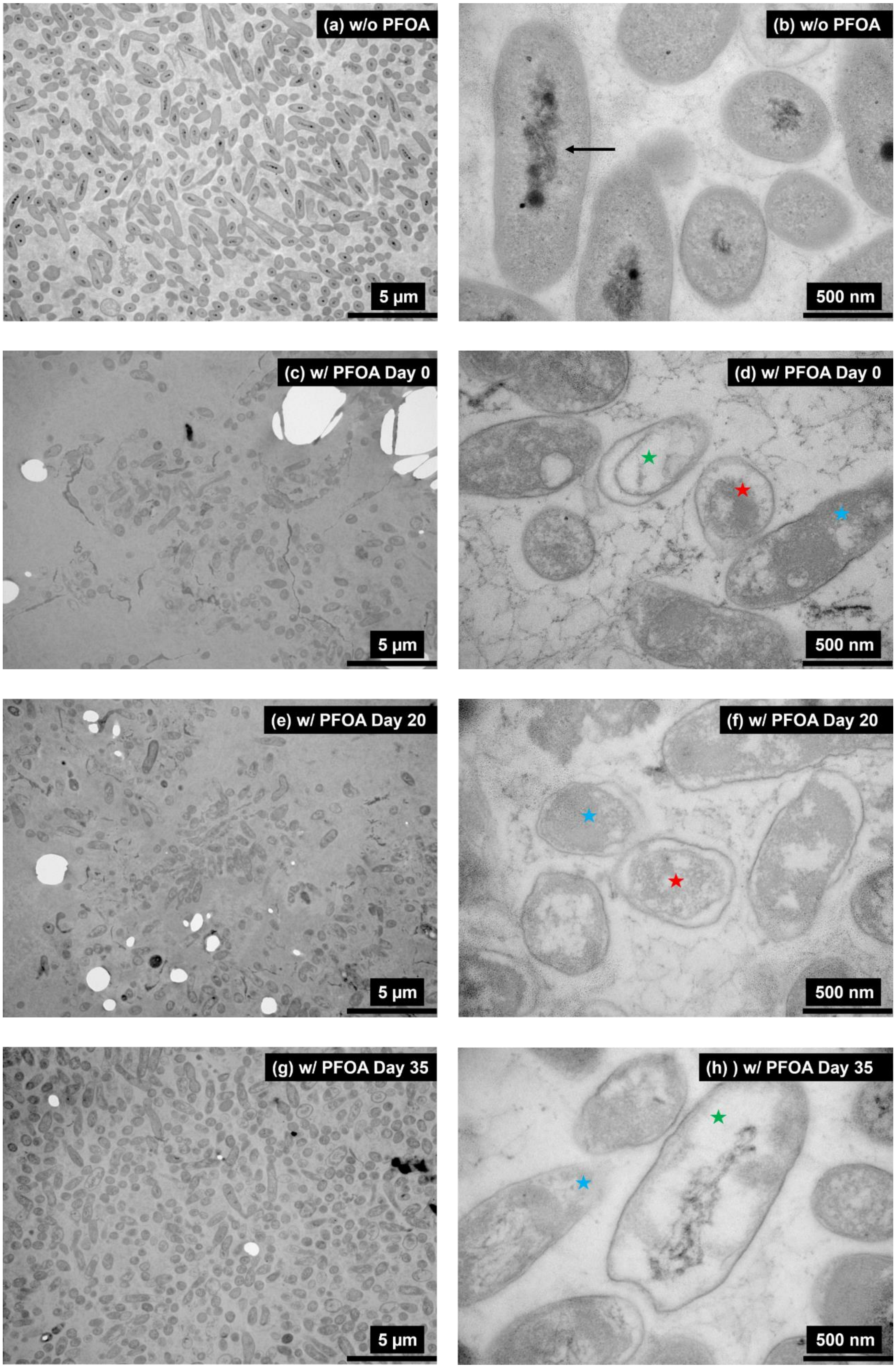

(a) w/o PFOA
5 μm
(b) w/o PFOA
500 nm
(c) w/ PFOA Day 0
5 μm
(d) w/ PFOA Day 0
500 nm
(e) w/ PFOA Day 20
5 μm
(f) w/ PFOA Day 20
500 nm
(g) w/ PFOA Day 35
5 μm
(h) ) w/ PFOA Day 35
500 nm

**Figure S2.5:** Wild type *R. palustris* without PFOA presence at (a) 5 µm and (b) 500 nm magnification. R palustris in PFOA spiked PM at day 0 (c and d), day 20 (e and f), and day 35 (g and h). The magnifications of images c, e, and g are at 5 µm, and images d, f, and h are at 500 nm. The microbe, in the absence of PFOA, is observed to be healthy (Black arrow indicating intact nucleus). However, most cells in (Figures S1d, S1f, and S1h) in PFOA presence have toxic effects (disrupted cell membranes (blue star), shrinkage (red star), lack of cytoplasm (green star) and nucleus) in the three samples from days 0, 20, and 35.
